# Declining autozygosity over time: an exploration in over 1 million individuals from three diverse cohorts

**DOI:** 10.1101/2022.10.13.512166

**Authors:** Sarah M.C. Colbert, Frank R. Wendt, Gita A. Pathak, Drew A. Helmer, Elizabeth R. Hauser, Matthew C. Keller, Renato Polimanti, Emma C. Johnson

## Abstract

We hypothesized that overall autozygosity is decreasing over generational time. In this report, we present data that partially support this hypothesis from three large cohorts of diverse ancestries, two from the US (All of Us and the Million Veteran Program, N=82,474 and 622,497, respectively) and one from the UK (UK Biobank, N=380,899). Our results from a mixed-effect meta-analysis demonstrate an overall trend of decreasing autozygosity over generational time (meta-analyzed slope=-0.029, se=0.009, p=6.03e-4). Using a chi-square difference test, we determined that a model including an ancestry-by-country interaction term fit the data best, indicating that ancestry differences in this trend differ by country. We found further evidence to suggest a difference between the US and UK cohorts by meta-analyzing within country, observing a significant negative estimate in the US cohorts (meta-analyzed slope=-0.058, se=0.015, p=1.50e-4) but a non-significant estimate in the UK (meta-analyzed slope=-0.001, se=0.008, p=0.945). We also found that the association between autozygosity and year of birth in the overall meta-analysis was substantially attenuated when accounting for educational attainment and income (meta-analyzed slope=-0.011, se=0.008, p=0.167), suggesting that increases in education and income may partially account for decreasing levels of autozygosity over time. To our knowledge, this is the largest demonstration of decreasing autozygosity over time in a modern sample (birth years 1904-2003), and we speculate that this trend can be attributed to increases in population size, urbanization and panmixia, with differences in demographic and sociocultural processes leading to country-specific differences in the rate of decline.

## MANUSCRIPT

There has been great interest in using measures of autozygosity - the proportion of the genome contained in runs of homozygosity (ROH) that are identical by descent (i.e., inherited from a common ancestor shared by both parents) - to examine evolutionary hypotheses about complex traits in humans ^1–3^ and to quantify the extent to which inbreeding depression impacts health and disease ^3–5^. While longer and more frequent ROHs are found in samples with close inbreeding, ROHs are ubiquitously found in samples across the world, even in seemingly outbred populations. By examining the proportion of the genome contained in ROHs (F_ROH_) alongside other measures of inbreeding (e.g., F_UNI_, the correlation between uniting gametes^6^), studies have shown how demographic history can influence the distribution of these different measures of inbreeding ^3,7^.

In a previous study^8^ using a sample of adolescents, we found an unexpectedly low mean level of autozygosity relative to previous autozygosity reports (mean F_ROH_ = 0.0005^8^ compared to 0.0016-0.007 ^9–11^) while the variance of F_ROH_ was similar to other studies. The particular sample used in that study, the Adolescent Brain Cognitive Development Study□ (ABCD Study®□)^12^, consisted of individuals who were much younger than most other samples analyzed in previous studies of autozygosity, with all individuals in the ABCD study having been born in 2006 or 2007. In researching this finding, we came across a study from Nalls *et al*. (2009)^13^, who found that in a sample of 809 North Americans of European descent aged 19-99 years old, autozygosity steadily declined in relation to birth year at a rate of 0.1% decrease in F_ROH_ for every 20 years decrease in chronological age. Aside from Nalls *et al*. (2009), there seem to be few mentions of this phenomenon in the literature, except for an interesting analysis of ancient DNA samples which found decreasing F_ROH_ over 1000s of years during the Holocene ^7^. We hypothesized that the relatively low level of autozygosity in the young ABCD Study sample might be reflective of secular trends of decreasing autozygosity over generational time in the modern era. In the previous study, we tested this by conducting a brief assessment of an independent cohort, the Collaborative Study on the Genetics of Alcoholism (COGA) ^14–16^, and observed a small but highly significant decrease in F_ROH_ with increasing birth year (standardized beta= −0.06, s.e.= 0.01, p= 2.5e-9)^8^. Based on this finding, we would predict a 0.001 decrease in F_ROH_ over a period of 100 years. However, this trend has so far only been examined in relatively small (N < 11,000) North American cohorts comprised mostly of individuals of European and African descent. Thus, it is unclear to what extent this association between F_ROH_ and birth year generalizes across different and more diverse samples.

In the current report, we sought to address this gap in the literature using data from three large cohorts spanning the US (All of Us (AoU), N = 82,474; Million Veteran Program (MVP), N = 622,497) and UK (UK Biobank (UKB), N = 380,899) which include individuals of six ancestry groups determined by genetic principal components, broadly defined as Admixed American ancestry (AMR), African ancestry (AFR), Central South Asian ancestry (CSA), East Asian ancestry (EAS), European ancestry (EUR), and Middle Eastern ancestry (MID).

As linkage disequilibrium patterns and allele frequencies can differ across genetic ancestry groups and potentially induce spurious associations due to population stratification, we performed ROH calling and F_ROH_ regressions separately in each genetic ancestry subset of the cohorts, before meta-analyzing to increase sample size and statistical power. Thus, initial association tests were conducted in unrelated individuals in each ancestry subset of each cohort using a linear fixed-effect regression model which tested for the effect of birth year on F_ROH_, controlling for age, sex, and the first 10 within-ancestry genetic principal components, as well as genotyping batch and assessment center in the UK Biobank (Table 1). In this report we avoid comparing the F_ROH_~birth year relationships between genetic ancestries because sample sizes in some genetic ancestry subsets are too small to draw substantive conclusions (but individual within-ancestry estimates of the F_ROH_~birth year association are presented in Figure 1b). Using the effect sizes from the ancestry- and cohort-specific models, we performed two separate meta-analyses. First, we meta-analyzed across all cohorts and genetic ancestry groups using a mixed-effect meta-analysis model. We first tested a model with main effects only (ancestry and country as fixed effects, cohort as a random effect); when we then tested a model with an interaction term between ancestry group and country, this model fit significantly better than the main effects-only model (chi-square difference = 27.156, p = 5.32e-5). Given this finding, we decided to also examine country-specific estimates; thus, we also present a mixed-effect meta-analysis (controlling for genetic ancestry group as a fixed effect and cohort as a random effect) of the two US cohorts and a fixed-effect meta-analysis of the UK cohort (since there was only one UK cohort, we did not need to include cohort as a random effect) to calculate and compare country-specific estimates. In this report, we present the meta-analyzed slope (beta_M) from our meta-analysis models; this represents the effect of birth year on F_ROH_ on average across ancestry groups, countries, and cohorts. We applied a Bonferroni correction to correct for six total tests: two models (main model, model correcting for educational attainment and income) meta-analyzed three ways (across all cohorts, only in US samples, only in UK samples), resulting in a significance threshold of p = 0.0083. We note that this threshold is somewhat conservative given the substantial overlap amongst the tests.

**Figure 1.**
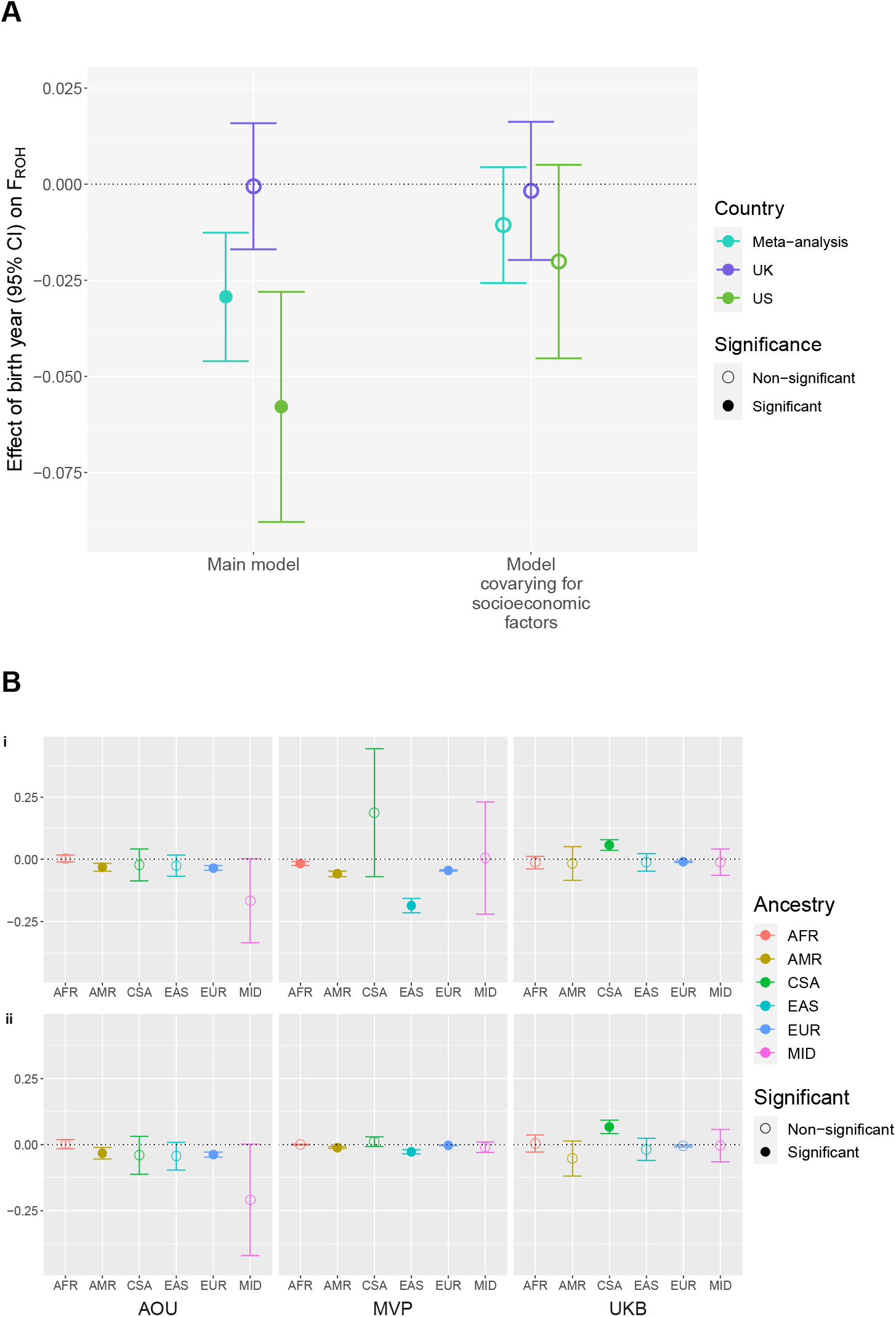
**(A)** Effect of birth year on F_ROH_ in each meta-analysis and model type. Points represent meta-analyzed slope values and bars represent 95% confidence intervals. Significance was determined using a conservative Bonferroni correction for 6 tests (3 types of meta-analysis [UK, US, and overall] and 2 possible models [main model, model controlling for socioeconomic factors]), resulting in a p-value threshold of 0.0083. **(B)** Effect of birth year on F_ROH_ in each ancestry and cohort. Points represent betas and bars represent 95% confidence intervals. Effects in the main model are shown in panel i, effects when controlling for educational attainment and income are shown in panel ii. Significance was determined using the previously mentioned threshold of p = 0.0083. AFR = African genetic ancestry; AMR = Admixed American genetic ancestry; CSA = Central South Asian genetic ancestry; EAS = East Asian genetic ancestry; EUR = European genetic ancestry; MID = Middle Eastern genetic ancestry.

In the primary meta-analysis across all ancestry groups and cohorts, birth year was negatively associated with F_ROH_ on average (beta_M = −0.029, se = 0.009, p= 6.03e-4; Figure 1a, Table S1). We found divergent effects in the within-country meta-analysis, observing a significant and strong negative effect of birth year on F_ROH_ in the US cohorts (beta_M = −0.058, se = 0.015, p = 1.50e-4), but a non-significant effect in the UK cohort (beta_M = −0.001, se = 0.008, p = 0.945). We note that a significant negative association was observed in the UKB sample of European descent (beta = −0.010, se = 0.002, p = 6.11e-9); still, the effect was much weaker than in the genetically-defined European ancestry subsets of the AoU (beta = −0.035, se = 0.005, p = 2.22e-13) and MVP (beta = −0.044, se = 0.001, p = 2.65e-195) cohorts (Figure 1b). This may reflect differences across the US and UK in terms of the rate of urbanization and/or demographic changes. While the percent of the population living in urban areas has surged 29% over the last 70 years in the US, urbanization in the UK has only increased by 6.2% ^17^, potentially contributing to the weaker changes in autozygosity in the UK cohort. Another possible reason for this difference is migration patterns; consistent immigration to the US from many different countries over the 20th century has facilitated more diverse and frequent admixture in Americans ^18^, leading to a more rapid decline in average autozygosity compared to the UK where immigration rates are lower ^19^. Furthermore, the physical isolation of Britain from the rest of Europe has presented challenges to migration historically ^20^, providing an explanation for the more stable rate of autozygosity in this population. We also acknowledge that the UK Biobank, compared to the two US cohorts, is much more limited in the chronological age span of its cohort. Individuals in the UK Biobank were born between 1936 and 1970, while individuals in the MVP and AoU cohorts had birth years ranging from 1904-1999 and 1915-2003, respectively. It is possible that the decline in autozygosity observed in the US cohorts may only become identifiable over many generations, as shorter periods of time may reflect short-term trends in response to historical and sociocultural changes.

We also estimated the association between birth year and a second measure of inbreeding, F_UNI_. Since F_ROH_ can better capture the effects of homozygosity at rare variants while F_UNI_ is thought to be a better measure of homozygosity at common variants ^3,5^, we tested both measures to determine whether common and/or rare variants were contributing to this trend in decreasing autozygosity, or whether variants contributing to this decline span a range of frequencies. We observed consistent direction of effects of F_UNI_ (albeit non-significant) in the overall meta-analysis (beta_M = −0.015, se = 0.009, p = 0.105) and the US-specific meta-analysis (beta_M = −0.045, se = 0.080, p = 0.011), while the estimate in the UK-specific meta-analysis was positive but non-significant (beta_M = 0.014, se = 0.008, p = 0.055). While the associations with F_UNI_ were weaker, the generally consistent direction of effects was unsurprising given the strong correlation between F_ROH_ and F_UNI_ measures (e.g., r^2^ in the genetically-defined European ancestry subset of AoU = 0.66) and suggests that these patterns of decline are not specific to F_ROH_ and likely represent trends in autozygosity more generally across variants of all frequencies. We note that, while imputed SNP data can lead to biased estimates of F_ROH_, estimates of F_UNI_ are more powerful and unbiased when derived from imputed data ^5^. Imputed data was not available for the AoU cohort and thus we have used non-imputed genotype array data to estimate both F_ROH_ and F_UNI_ in all cohorts for consistency, acknowledging that our estimates of F_UNI_ may be underpowered.

Previous studies have demonstrated strong relationships between educational attainment, social mobility, and autozygosity, with greater educational attainment correlating with more mobility^21^, and greater mobility in the parental generation mediating observed relationships between their educational attainment and their child’s autozygosity^11^. To investigate whether differences in educational attainment and other socioeconomic factors such as income might be responsible for the observed decline in autozygosity over time, we tested an additional model in which birth year, educational attainment and income simultaneously predicted F_ROH_ (while controlling for the same covariates as above, see Supplemental Material and Methods). After meta-analyzing across cohorts and genetic ancestry groups, the effect of birth year on F_ROH_ was attenuated (beta_M = −0.011, se = 0.008, p = 0.167; Figure 1a) when educational attainment and income were included in the model. We subsequently meta-analyzed within countries and found that educational attainment and income substantially weakened the effect in the US cohorts (beta_M = −0.020, se = 0.013, p = 0.117) (Figure 1a). In the UK, where the association between F_ROH_ and birth year was already close to null when averaged across ancestry groups, controlling for educational attainment and income had no notable effect on the F_ROH_~birth year relationship (beta_M = −0.002, se = 0.009, p = 0.848). We speculated that generations have become increasingly more educated over time and this has changed patterns in mobility; perhaps these patterns of increased geographic mobility, acting in concert with assortative mating on socioeconomic status, have partially contributed to the observed decrease in autozygosity over time. To test whether levels of education and income have increased over generations, we regressed educational attainment and income on birth year and indeed observed a significant increase in educational attainment over time (beta_M = 0.102, se = 0.034, p = 0.003), while the change in income was not significant (beta_M = −0.081, se = 0.116, p = 0.487). Within-country meta-analyses revealed a much stronger positive relationship between educational attainment and birth year in the UK (beta_M = 0.136, se = 0.009, p = 1.04e-56) than in the US (beta_M = 0.069, se = 0.068, p = 0.309). Furthermore, this null result in the US meta-analysis seemed to be driven by conflicting ancestry-specific results in the AoU cohort, with the two largest ancestry groups showing significant *negative* relationships between educational attainment and birth year, and the third-largest ancestry group demonstrating a significant association in the expected, positive direction (Supplemental Table S1). Results did not change when we restricted the age range in AoU to match the birth years of the UK Biobank (1936-1970).

Like Nalls *et al*. (2009), we consider that the overall pattern of decreasing autozygosity may be associated with population growth, urbanization, and increased mobility. Population sizes have increased both in the US and worldwide ^22^ and previous analyses have noted that rapid growth in population size or large effective population size is associated with a decrease in autozygosity ^3,7,23^. For example, a study from Ceballos *et al*. (2021) found a decrease in F_ROH_ over 1000s of years during the Holocene, likely in response to population growth arising from the development of agriculture at the time. Population expansion, therefore, appears to contribute to decreases in autozygosity over both short and long time periods, as well as in both modern and ancient samples. In addition to modern population growth, the flocking of individuals from many small, isolated rural areas to densely populated cities breaks down previous geographic and population barriers to panmixia, in turn reducing endogamy and increasing the likelihood that individuals mate with those who are more genetically different from themselves ^24^. Our results also suggest that socioeconomic factors, especially educational attainment, at least partially explain the F_ROH_~birth year relationship. We found that educational attainment is higher on average in more recent generations, although this relationship was stronger in the UK Biobank and the MVP cohorts than in AoU, where results were mixed. One previous study found that those with higher educational attainment were more likely to move large distances away from their birthplace and subsequently mate with an individual that is less closely related to them but who also shares a similarly high level of educational attainment. As a result, offspring of these individuals were more outbred (had low levels of autozygosity) and would have inherited genes associated with greater educational attainment^11^. As individuals became increasingly more educated, this pattern of migration and mating may have become more common, leading to overall declines in average autozygosity. It may also be that increased globalization and mobility are reflected in higher levels of educational attainment ^25,26^, which then are associated with lower autozygosity on average in the countries we have studied. Still, the relationships between socioeconomic factors and birth year were not as clear-cut in the US cohorts as in the UK Biobank, and further studies are needed to clarify the role of these factors in the observed decline in autozygosity.

Nalls *et al*. (2009) also hypothesized that decreasing autozygosity should correlate with decreasing rates of rare recessive genetic diseases, while Campbell *et al*. (2009)^27^ estimated that this effect measured by Nalls *et al*. (2009) has prevented 1% of the annual births that would be affected with an autosomal recessive disorder. We might also expect slight changes in complex traits that are partly influenced by recessive variants, such as cognitive abilities. We used our estimated rates of declining autozygosity and estimates of associations between F_ROH_ and complex traits from published literature^3^ to predict estimated changes in several traits. For example, based on our findings in the European-ancestry subset of the AoU sample and published associations in Clark *et al*. (2019), we predict a 0.004 standard deviation increase in cognitive *g*, a 0.019 kg increase in grip strength, a 0.019 cm increase in height, and a 0.0095 year increase in educational attainment over a 100-year period due to decreases in autozygosity. Of course, these are only illustrative predictions, but we expect that while declining autozygosity might have small effects on complex traits, such as those estimated here, this decline may show more appreciable effects on traits and diseases that are more strongly influenced by rare, recessive genetic variants.

Importantly, we also note that these findings shed light on the consequences of overlooking sample composition - including range of birth years - when conducting comparisons of inbreeding across populations. Future studies that wish to analyze measures of inbreeding, such as F_ROH_, across populations should be aware that sample differences not only in geography or genetically-defined ancestry groups, but also in age, can affect the mean level of F_ROH_.

We note several limitations to the current study, the first being that our analyses only include samples from the US and UK. Given the differences observed between the US and UK cohorts, we would also expect changes in autozygosity over time to differ in cohorts from other countries in response to region-specific cultural practices (e.g., consanguinity) and demographic trends (e.g., migration rates). As biobanks in other countries continue to grow and include more diverse samples, we will be better able to assess how this pattern may differ from country to country. While we were able to include a diverse sample encompassing individuals from six different genetic ancestry clusters, a major limitation of our sample (N = 1,085,870) is that it still consisted mainly of individuals with European genetic ancestry (N = 847,427; 78.0%). Therefore, the overall generalizability of our findings across samples of non-European ancestry groups is limited. Furthermore, the degree of admixture in individuals in this study likely varies amongst the different genetic ancestry groups and cohorts. For example, a majority of the individuals in the genetically defined American and African ancestry subsets of the UK Biobank are likely admixed and share ancestry with the individuals in the European ancestry subset. On the other hand, individuals in the UK Biobank with less common patterns of admixture could not be grouped into sufficient sized groups and were thus excluded by the PanUKB analysis team^28^. Cross-ancestry mating is likely a contributing factor to declining autozygosity, and by excluding some individuals with admixture we are likely under-estimating the true decline in autozygosity over time. Finally, while we show that educational attainment and income partly drive the observed association, we were unable to investigate how other variables linked to assortative mating, such as religiosity, may also influence autozygosity ^29^.

In summary, we demonstrate an overall trend of declining autozygosity over time on average across multiple ancestry groups and countries, with a stronger overall effect in the US than in the UK. Controlling for educational attainment and income substantially attenuates this relationship but does not fully explain the decline in autozygosity observed. We hypothesize that population growth combined with increased urbanization, globalization, and mobility are likely to be driving this trend. Future research should assess the relationship between autozygosity and birth year in better-powered samples of more diverse ancestry groups and ages in order to determine how autozygosity has changed across different time spans and regions.

## SUPPLEMENTAL MATERIAL AND METHODS

### Samples

This study used data from two North American samples, the All of Us biobank (AoU) and the Million Veteran Program (MVP). We stratified these cohorts into six categorical ancestry groups commonly defined by genetic principal components: African, Admixed American, Central/South Asian, East Asian, European, and Middle Eastern. These were defined using reference populations from the 1000 Genomes Project^30^ and the Human Genome Diversity Project^31^ as previously reported^32^.

The AoU research program includes over 1 million diverse individuals from across the U.S. and combines data from a variety of sources, such as an initial physical examination, follow-up self-report surveys, electronic health records and even genetic data from a subset of individuals. Individuals with genetic data spanned a wide range of chronological ages (birth years between 1915-2003), making the sample ideal for the current study. While our analyses use array data, the AoU biobank only provides ancestry assignments and relatedness data for the whole-genome sequencing dataset (N = 98,590), leaving us with 82,474 unrelated, genotyped individuals with ancestry assignments to use in the current analysis. We opted to use unrelated individuals in our analyses as the power that would have been gained by including related individuals would have been relatively small whereas the increase in computational resources and time required to control for relatedness in our analyses would be large and likely exceed the resources available to us via the All of Us Researcher workbench. For similar reasons we used the pre-computed ancestry assignments from the AoU dataset.

The MVP sample also includes individuals with a wide range of birth years (1904-1999) and is highly diverse. A detailed description of ancestry prediction in the MVP sample has been discussed previously^32^. We used KING^33^ to identify pairs of individuals who were estimated to be third-degree relatives or closer (kinship coefficient > 0.0442) and randomly removed one individual from each pair. Restricting to unrelated individuals left 622,497 individuals from MVP in our analyses.

To assess how trends of changing autozygosity over time may differ across countries, we also included data from the UKB (N ~ 500,000), which has collected genetic samples from individuals born between 1936 and 1970 at 23 assessment centers across the United Kingdom. We used ancestry and relatedness assignments provided by the Pan-UKB Team^28^ to remove related individuals (N = 65,887) and subset the UKB sample into the six previously mentioned genetically predicted ancestry categories. Doing so resulted in a total of 380,899 unrelated individuals.

### Measures

#### Educational Attainment

Educational attainment data in the AoU sample was collected by asking individuals the highest grade or year of school they completed (item concept = educationlevel_highestgrade) and individuals were given eight choices (e.g., “never attended school or only attended kindergarten”, “grades 1 through 4 (Primary school)”, etc.) to choose from. Choices were equated to approximate “years of education”, averaged across the possible range of a category, such that, for example, an individual who selected “grades 1 through 4” would be assigned 2.5 years of education. The UKB dataset does not provide a “years of education” measure, but does record each individual’s educational qualifications (Data-Field 6138). Qualifications in the UKB dataset were mapped to the International Standard Classification of Education (ISCED) levels and then converted to a “years of education” value, following the procedure from Okbay *et al*. (2016)^34^. Individuals with multiple qualifications were assigned a “years of education” value corresponding to the highest qualification. In the MVP dataset, educational attainment was measured using 7 answers (lowest being “less than high school’ and highest being “professional or doctorate degree”’) to the question “What is the highest degree or level of school you have completed?” which were then recoded into numeric values 1-7.

#### Income

The AoU dataset provided annual household income (item concept = income_annualincome) in the form of an ordinal measure with 9 categories (e.g., “less than $10,000”, “$10,000-24,999”, etc.), which we re-assigned to corresponding numeric values 1-9. Similarly, UKB annual household income (Data-Field 738) data was also in the form of an ordinal variable, and we transformed the five income brackets into numeric values 1-5. Finally, in MVP, annual household income was also reported as an ordinal variable, with ten income brackets being recoded to numeric values 1-10.

### Analyses

We performed F_ROH_ and F_UNI_ estimation and association testing separately for each ancestry group within each cohort. Following the procedure from Clark *et al*. (2019), we used PLINK 1.9^35^ to clean the genotypic data before calling ROHs and estimating F_ROH_ and F_UNI_. Genotypic data cleaning consisted of excluding SNPs with > 3% missingness or MAF < 5% and individuals with > 3% missing data. The resulting data was used to call ROHs in PLINK 1.9, using the following parameters: --homozyg-window-snp 50; -- homozyg-snp 50; --homozyg-kb 1500; --homozyg-gap 1000; --homozyg-density 50; -- homozyg-window-missing 5; homozyg-window-het 1. No linkage disequilibrium pruning was performed. We calculated F_ROH_ as the total length of ROHs summed for each individual, and then divided by the total SNP-mappable autosomal distance (2.77 × 10^6^ kilobases). F_UNI_ was estimated using the --ibc command in PLINK 1.9 (F_UNI_ corresponding to ‘Fhat3’ in the output, the correlation between uniting gametes^6^).

We performed multiple linear regression models to determine if there was a significant effect of birth year on autozygosity in our samples. In the UKB sample, fixed-effect regression models controlled for sex, genotyping batch, assessment center and the first 10 genetic within ancestry principal components as covariates. In the MVP and AoU cohorts, we used fixed-effect regression models to control for sex and the first 10 genetic within ancestry principal components as covariates.

In addition to these main models, we conducted a follow-up analysis in which we tested for a mediating effect of socioeconomic status by constructing models in which we covaried for educational attainment and income (along with the original covariates mentioned above). Separately, we measured trends in these socioeconomic factors over time by regressing them on birth year (e.g., educational attainment ~ birth year). In these models we only covaried for the non-genetic covariates listed above.

All models were run separately by genetic ancestry and cohort. Meta-analyses were performed in R using the metafor package ^36^. To meta-analyze across all cohorts and ancestries, we used a mixed-effect meta-analysis model, controlling for ancestry and country as fixed effects, an interaction term between ancestry and country, and cohort as a random effect (e.g., rma.mv(yi=estimate, V=sampvar, mods = ~ancestry.c*country.c, random = ~1|cohort, data=dat, method=“ML”). We chose to include the interaction term between ancestry and country after using a chi-squared difference test to compare the fit between the model including this interaction and the model without the interaction. To be consistent, meta-analyses of results from all other models (e.g., models controlling for educational attainment and income) also include the interaction term.

## Acknowledgements

The authors thank Keith Lohse for his help implementing and interpreting mixed-effect meta-analysis models. Analyses were performed under UKB application #58146.

The All of Us Research Program is supported by the National Institutes of Health, Office of the Director: Regional Medical Centers: 1 OT2 OD026549; 1 OT2 OD026554; 1 OT2 OD026557; 1 OT2 OD026556; 1 OT2 OD026550; 1 OT2 OD 026552; 1 OT2 OD026553; 1 OT2 OD026548; 1 OT2 OD026551; 1 OT2 OD026555; IAA #: AOD 16037; Federally Qualified Health Centers: HHSN 263201600085U; Data and Research Center: 5 U2C OD023196; Biobank: 1 U24 OD023121; The Participant Center: U24 OD023176; Participant Technology Systems Center: 1 U24 OD023163; Communications and Engagement: 3 OT2 OD023205; 3 OT2 OD023206; and Community Partners: 1 OT2 OD025277; 3 OT2 OD025315; 1 OT2 OD025337; 1 OT2 OD025276. In addition, the All of Us Research Program would not be possible without the partnership of its participants.

## Data and code availability

The UK Biobank data used in this study are available from the UK Biobank by applying for access via the Access Management System (https://www.ukbiobank.ac.uk/enable-your-research/apply-for-access). Original All of Us Biobank data are available to registered and approved All of Us researchers (https://www.researchallofus.org/register/). Genetic data requires controlled tier access, which researchers can register for through their institutions. Data from the Million Veteran Program are only available to VA investigators and other approved partners.

Code to analyze the UK Biobank data is available via github: https://github.com/sarahcolbert/autozygosity_time_ukbb. For privacy reasons, we are unable to share the code used to analyze the AoU and MVP data, but the code is almost identical to the code used to analyze the UK Biobank data.

---

**Table S1.**
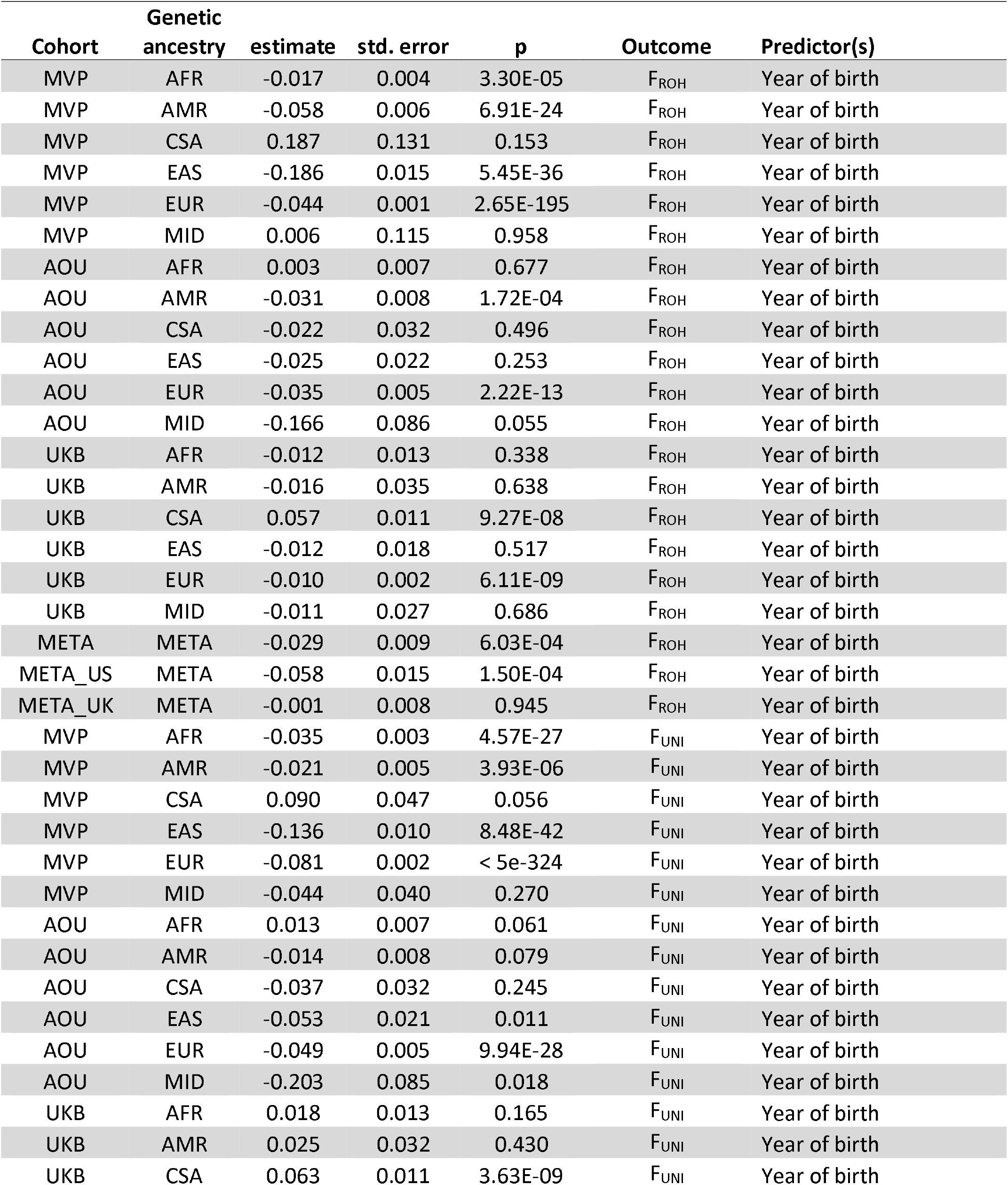
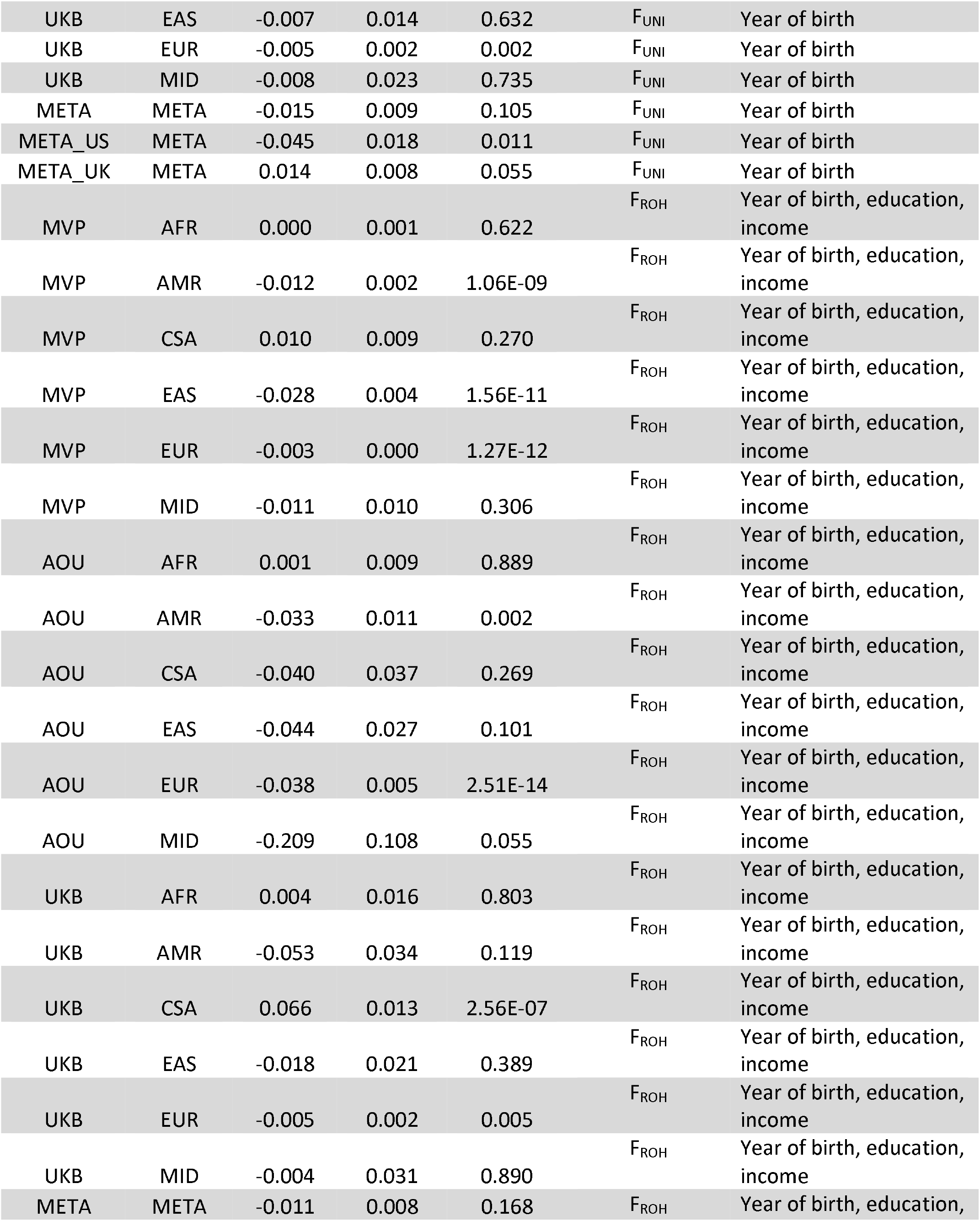
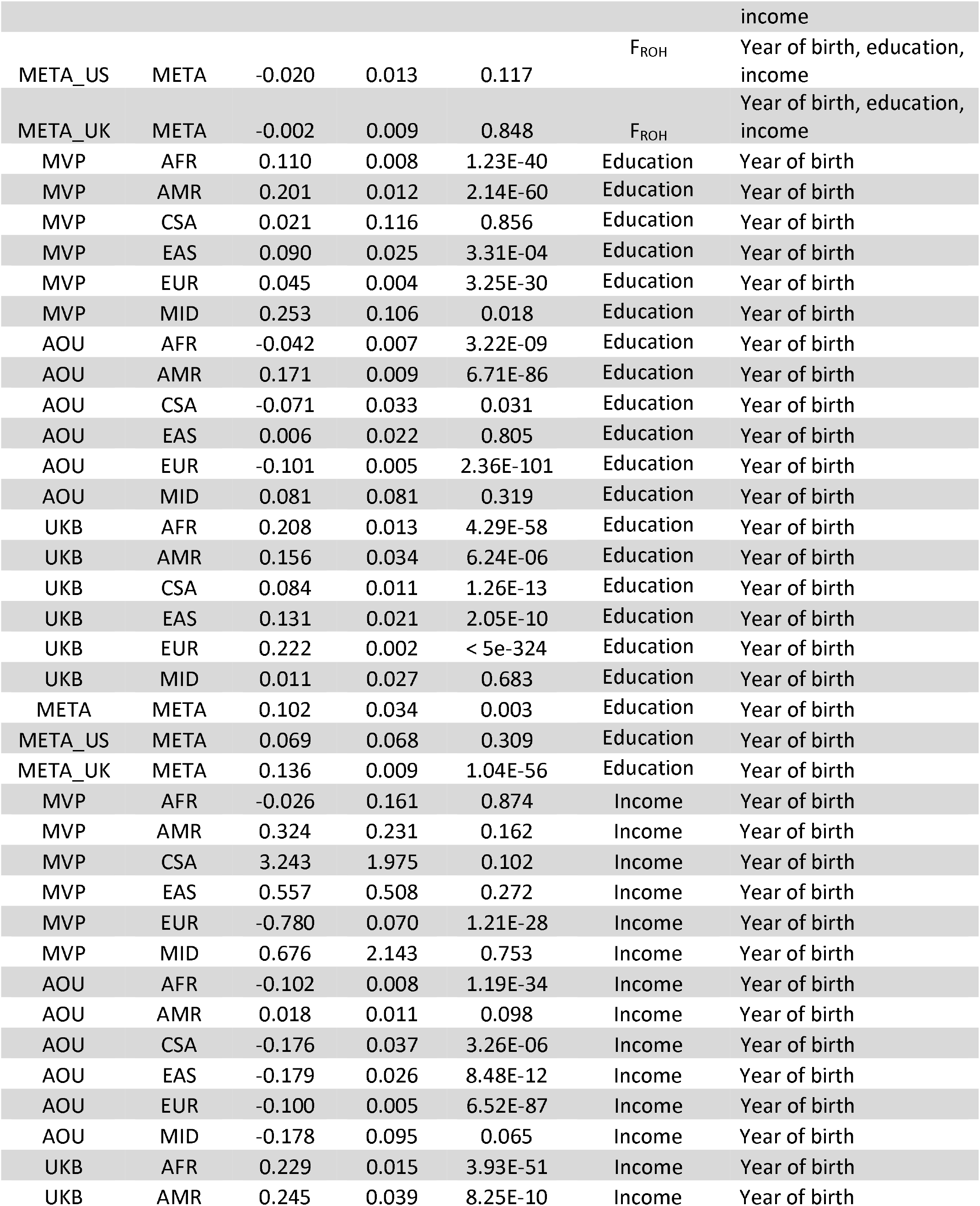
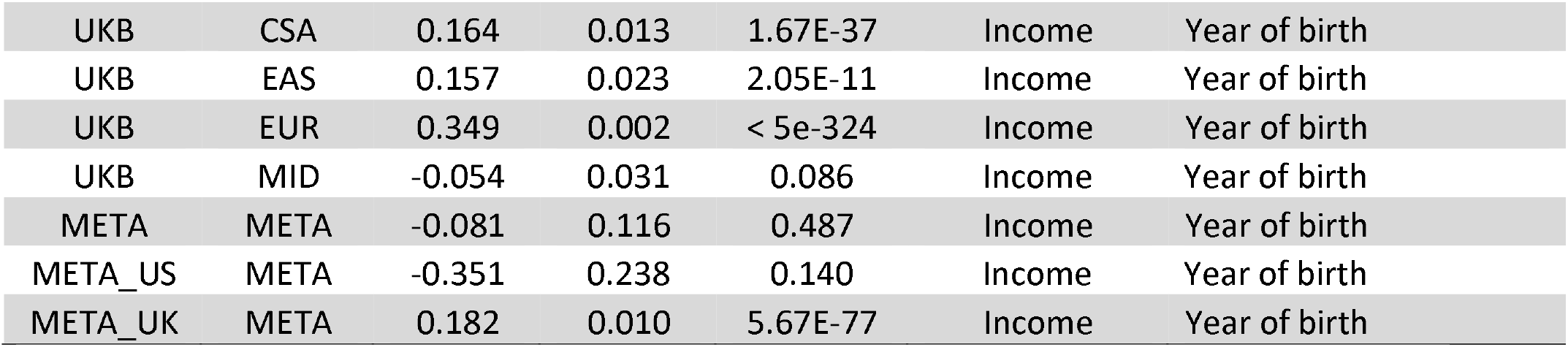
Results from all models.

## References

1. Yengo, L., Zhu, Z., Wray, N.R., Weir, B.S., Yang, J., Robinson, M.R., and Visscher, P.M. (2017). Detection and quantification of inbreeding depression for complex traits from SNP data. Proc Natl Acad Sci U S A 114. 10.1073/pnas.1621096114.

2. Johnson, E.C., Evans, L.M., and Keller, M.C. (2018). Relationships between estimated autozygosity and complex traits in the UK Biobank. PLoS Genet 14. 10.1371/journal.pgen.1007556.

3. Clark, D.W., Okada, Y., Moore, K.H.S., Mason, D., Pirastu, N., Gandin, I., Mattsson, H., Barnes, C.L.K., Lin, K., Zhao, J.H., et al. (2019). Associations of autozygosity with a broad range of human phenotypes. Nat Commun 10. 10.1038/s41467-019-12283-6.

4. Ceballos, F.C., Hazelhurst, S., Clark, D.W., Agongo, G., Asiki, G., Boua, P.R., Xavier Gómez-Olivé, F., Mashinya, F., Norris, S., Wilson, J.F., et al. (2020). Autozygosity influences cardiometabolic disease-associated traits in the AWI-Gen sub-Saharan African study. Nature Communications 2020 11:1 11, 1–8. 10.1038/s41467-020-19595-y.

5. Yengo, L., Yang, J., Keller, M.C., Goddard, M.E., Wray, N.R., and Visscher, P.M. (2021). Genomic partitioning of inbreeding depression in humans. The American Journal of Human Genetics 108, 1488–1501. 10.1016/J.AJHG.2021.06.005.

6. Yang, J., Lee, S.H., Goddard, M.E., and Visscher, P.M. (2011). GCTA: A Tool for Genome-wide Complex Trait Analysis. Am J Hum Genet 88, 76. 10.1016/J.AJHG.2010.11.011.

7. Ceballos, F.C., Gürün, K., Altinişik, N.E., Gemici, H.C., Karamurat, C., Koptekin, D., Vural, K.B., Mapelli, I., Saĝlican, E., Sürer, E., et al. (2021). Human inbreeding has decreased in time through the Holocene. Current Biology 31. 10.1016/j.cub.2021.06.027.

8. Colbert, S.M., Keller, M.C., Agrawal, A., and Johnson, E.C. (2022). Exploring the Relationships Between Autozygosity, Educational Attainment, and Cognitive Ability in a Contemporary, Trans-Ancestral American Sample. Behavior Genetics 2022, 1–9. 10.1007/S10519-022-10113-Y.

9. Power, R.A., Nagoshi, C., DeFries, J.C., and Plomin, R. (2014). Genome-wide estimates of inbreeding in unrelated individuals and their association with cognitive ability. European Journal of Human Genetics 22. 10.1038/ejhg.2013.155.

10. Howrigan, D.P., Simonson, M.A., Davies, G., Harris, S.E., Tenesa, A., Starr, J.M., Liewald, D.C., Deary, I.J., McRae, A., Wright, M.J., et al. (2016). Genome-wide autozygosity is associated with lower general cognitive ability. Mol Psychiatry 21. 10.1038/mp.2015.120.

11. Abdellaoui, A., Hottenga, J.J., Willemsen, G., Bartels, M., van Beijsterveldt, T., Ehli, E.A., Davies, G.E., Brooks, A., Sullivan, P.F., Penninx, B.W.J.H., et al. (2015). Educational Attainment Influences Levels of Homozygosity through Migration and Assortative Mating. PLoS One 10, e0118935. 10.1371/JOURNAL.PONE.0118935.

12. Jernigan, T.L., Brown, S.A., and Dowling, G.J. (2018). The Adolescent Brain Cognitive Development Study. Journal of Research on Adolescence 28. 10.1111/jora.12374.

13. Nalls, M.A., Simon-Sanchez, J., Gibbs, J.R., Paisan-Ruiz, C., Bras, J.T., Tanaka, T., Matarin, M., Scholz, S., Weitz, C., Harris, T.B., et al. (2009). Measures of autozygosity in decline: Globalization, urbanization, and its implications for medical genetics. PLoS Genet 5. 10.1371/journal.pgen.1000415.

14. Begleiter, H., Reich, T., Hesselbrock, V., Porjesz, B., Li, T.-K., Schuckit, M.A., Edenberg, H.J., and Rice, J.P. (1995). The collaborative study on the genetics of alcoholism: An update. Alcohol Research and Health 19, 228–236.

15. Bucholz, K.K., McCutcheon, V. V., Agrawal, A., Dick, D.M., Hesselbrock, V.M., Kramer, J.R., Kuperman, S., Nurnberger, J.I., Salvatore, J.E., Schuckit, M.A., et al. (2017). Comparison of Parent, Peer, Psychiatric, and Cannabis Use Influences Across Stages of Offspring Alcohol Involvement: Evidence from the COGA Prospective Study. Alcohol Clin Exp Res 41, 359–368. 10.1111/ACER.13293.

16. Nurnberger, J.I., Wiegand, R., Bucholz, K., O’Connor, S., Meyer, E.T., Reich, T., Rice, J., Schuckit, M., King, L., Petti, T., et al. (2004). A family study of alcohol dependence: coaggregation of multiple disorders in relatives of alcohol-dependent probands. Arch Gen Psychiatry 61, 1246–1256. 10.1001/ARCHPSYC.61.12.1246.

17. UNDESA (2018). World Urbanization Prospects□: The 2018 Revision.

18. Nothnagel, M., Tehua Lu, T., Kayser, M., and Krawczak, M. (2010). Genomic and geographic distribution of SNP-defined runs of homozygosity in Europeans. Hum Mol Genet 19, 2927–2935. 10.1093/hmg/ddq198.

19. United Nations (2019). International migration 2019 report.

20. O’Dushlaine, C.T., Morris, D., Moskvina, V., Kirov, G., Gill, M., Corvin, A., Wilson, J.F., and Cavalleri, G.L. (2010). Population structure and genome-wide patterns of variation in Ireland and Britain. European Journal of Human Genetics 18, 1248. 10.1038/EJHG.2010.87.

21. Abdellaoui, A., Hugh-Jones, D., Yengo, L., Kemper, K.E., Nivard, M.G., Veul, L., Holtz, Y., Zietsch, B.P., Frayling, T.M., Wray, N.R., et al. (2019). Genetic correlates of social stratification in Great Britain. Nature Human Behaviour 2019 3:12 3, 1332–1342. 10.1038/s41562-019-0757-5.

22. Roser, M., Ritchie, H., and Ortiz-Ospina, E. (2020). World Population Growth-Our World in Data. Population Reference Bureau.

23. Keller, M.C., Visscher, P.M., and Goddard, M.E. (2011). Quantification of inbreeding due to distant ancestors and its detection using dense single nucleotide polymorphism data. Genetics 189. 10.1534/genetics.111.130922.

24. Rudan, I., Carothers, A.D., Polasek, O., Hayward, C., Vitart, V., Biloglav, Z., Kolcic, I., Zgaga, L., Ivankovic, D., Vorko-Jovic, A., et al. (2008). Quantifying the increase in average human heterozygosity due to urbanisation. European Journal of Human Genetics 16. 10.1038/ejhg.2008.48.

25. Meyer, J.W. (2007). Globalization: Theory and Trends. http://dx.doi.org/10.1177/0020715207079529 48, 261–273. 10.1177/0020715207079529.

26. Schofer, E., and Meyer, J.W. (2005). The Worldwide Expansion of Higher Education in the Twentieth Century. Source 70, 898–920.

27. Campbell, H., Rudan, I., Bittles, A.H., and Wright, A.F. (2009). Human population structure, genome autozygosity and human health. Genome Med 1. 10.1186/gm91.

28. Pan-UKB team (2020). https://pan.ukbb.broadinstitute.org.

29. Abdellaoui, A., Hottenga, J.J., Xiao, X., Scheet, P., Ehli, E.A., Davies, G.E., Hudziak, J.J., Smit, D.J.A., Bartels, M., Willemsen, G., et al. (2013). Association between Autozygosity and Major Depression: Stratification due to Religious Assortment. Behav Genet 43, 455–467. 10.1007/S10519-013-9610-1.

30. Auton, A., Abecasis, G.R., Altshuler, D.M., Durbin, R.M., Bentley, D.R., Chakravarti, A., Clark, A.G., Donnelly, P., Eichler, E.E., Flicek, P., et al. (2015). A global reference for human genetic variation. Nature 526. 10.1038/nature15393.

31. Li, J.Z., Absher, D.M., Tang, H., Southwick, A.M., Casto, A.M., Ramachandran, S., Cann, H.M., Barsh, G.S., Feldman, M., Cavalli-Sforza, L.L., et al. (2008). Worldwide human relationships inferred from genome-wide patterns of variation. Science (1979) 319, 1100–1104. 10.1126/SCIENCE.1153717/SUPPL_FILE/LI_SOM.PDF.

32. Wendt, F.R., Pathak, G.A., Vahey, J., Qin, X., Koller, D., Cabrera-Mendoza, B., Haeny, A., Harrington, K.M., Rajeevan, N., Duong, L.M., et al. (2022). Modeling the longitudinal changes of ancestry diversity in the Million Veteran Program. bioRxiv, 2022.01.24.477583. 10.1101/2022.01.24.477583.

33. Manichaikul, A., Mychaleckyj, J.C., Rich, S.S., Daly, K., Sale, M., and Chen, W.M. (2010). Robust relationship inference in genome-wide association studies. Bioinformatics 26, 2867. 10.1093/BIOINFORMATICS/BTQ559.

34. Okbay, A., Beauchamp, J.P., Fontana, M.A., Lee, J.J., Pers, T.H., Rietveld, C.A., Turley, P., Chen, G.B., Emilsson, V., Meddens, S.F.W., et al. (2016). Genome-wide association study identifies 74 loci associated with educational attainment. Nature 2016 533:7604 533, 539–542. 10.1038/nature17671.

35. Chang, C.C., Chow, C.C., Tellier, L.C.A.M., Vattikuti, S., Purcell, S.M., and Lee, J.J. (2015). Second-generation PLINK: Rising to the challenge of larger and richer datasets. Gigascience 4, s13742–015. 10.1186/s13742-015-0047-8.

36. Viechtbauer, W. (2010). Conducting Meta-Analyses in R with the metafor Package. J Stat Softw 36, 1–48. 10.18637/JSS.V036.I03.

